# Non-Electrostatic Basis for an Artificial Metalloenzyme Catalysis

**DOI:** 10.1101/2024.02.27.582238

**Authors:** Anagh Mukherjee, Subhendu Roy

## Abstract

The artificial metalloenzyme (referred to as **Ir-Q**) reported by Hartwig and coworkers presented an important milestone in merging the extraordinary efficiency of biocatalyst with the versatility of small molecule chemical catalyst in catalyzing a new-to-nature carbene insertion reaction. The artificial enzyme results from formal replacement of the Fe by an Ir(Me) moiety along with four C317G, T213G, L69V, V254L mutations in a natural Cytochrome enzyme CYP119 by directed evolution method. Importantly, this is a show-stopper enzyme as it exhibits a catalytic rate enhancement similar to that of natural enzymes. Despite this remarkable discovery, there is no mechanistic understanding as to why it displays extraordinary efficiency, so far been intractable to experimental methods. In this study, we have deciphered the ‘catalytically active conformation’ of **Ir-Q** using large-scale molecular dynamics simulations and rigorous quantum chemical calculations. Our study reveals how directed evolution mutations precisely position the cofactor-substrate in an unusual orientation within a reshaped active site that emerged during evolution and fostered by C−H…π interactions from more ordered mutated L69V and V254L residues. This productive conformation correctly reproduces the experimental barrier height and the catalytic effect of 2.7 kcal/mol, in excellent agreement with observed rate enhancement. Moreover, the active conformation features an unprecedented bonding interaction in a metal-carbene species that preferentially stabilizes the rate determining formation of an Iridium-Porphyrin Carbene intermediate to render the observed high catalytic rate acceleration. While the electrostatic criteria are widely established, this study suggests a new design paradigm towards realization of fully programmable protein catalysis.

## Introduction

Enzymes are exceptionally powerful biocatalysts which play crucial roles in nearly all biochemical processes.(1)The enormous rate accelerations and unparallel selectivity achievable with enzymes make them attractive catalysts for solving problems to humankind, including the energy crisis, pharmaceuticals sector, environmental pollution, and food shortages.(2, 3)Moreover, the functional capabilities of natural enzymes can be greatly expanded to accelerate chemical transformations by the artificial introduction of metal ion cofactors that facilitate redox chemistry, radical processes and challenging chemical reactions.(4)However, proper placement of protein, substrate and transition metal complex with a productive interaction has proven challenging, and consequently the catalytic efficiencies attained by these metalloenzymes are often lower than the isolated small-molecule complex.(5, 6)

In this view, Hartwig and coworkers have achieved an important milestone by reporting an artificial metalloenzyme [Ir(Me)-PIX-CYP119-Max, hereafter referred to as **Ir-Q**], derived from formal replacement of the Fe by an Ir(Me) moiety along with incorporating four active site C317G, T213G, L69V, V254L mutations in a natural Cytochrome P450 enzyme CYP119 by directed evolution method (Fig. 1a).(7)**Ir-Q** is a show-stopper enzyme as it shows surprisingly a similar kinetics with native enzymes by catalyzing new-to-nature reactions of intra- and intermolecular carbene insertion into the unactivated C−H bonds to produce important chemicals (Fig. 1b). Moreover, it merges the extraordinary efficiency and selectivity of a biological catalyst with the versatility of small molecule chemical catalyst to create highly active, productive, and selective metalloenzymes. However, the origin of the extraordinary efficiency of the most efficient mutant, **Ir-Q** is still unknown, which has so far been intractable to experimental methods. Understanding the underlying physical principles and reaction mechanism of artificial enzymes generated by directed evolution methods is fundamentally important to decipher protein structure-function correlation.(3) Recently, Hartwig and coworkers has reported crystal structure and crucial mechanistic details on a closely related, artificial metalloenzyme [Ir(Me)-PIX-CYP119, C317G, T213G, V254L, L155W, hereafter as **Ir-ArM**).(8) Using kinetic and other studies, they demonstrated that the Iridium Porphyrin Carbene (**Ir-PC**) intermediate formation step is indeed the rate determining step (RDS) in this **Ir-ArM**, which is almost similar to our target metalloenzyme differing only in a single mutation.

**Fig. 1.**
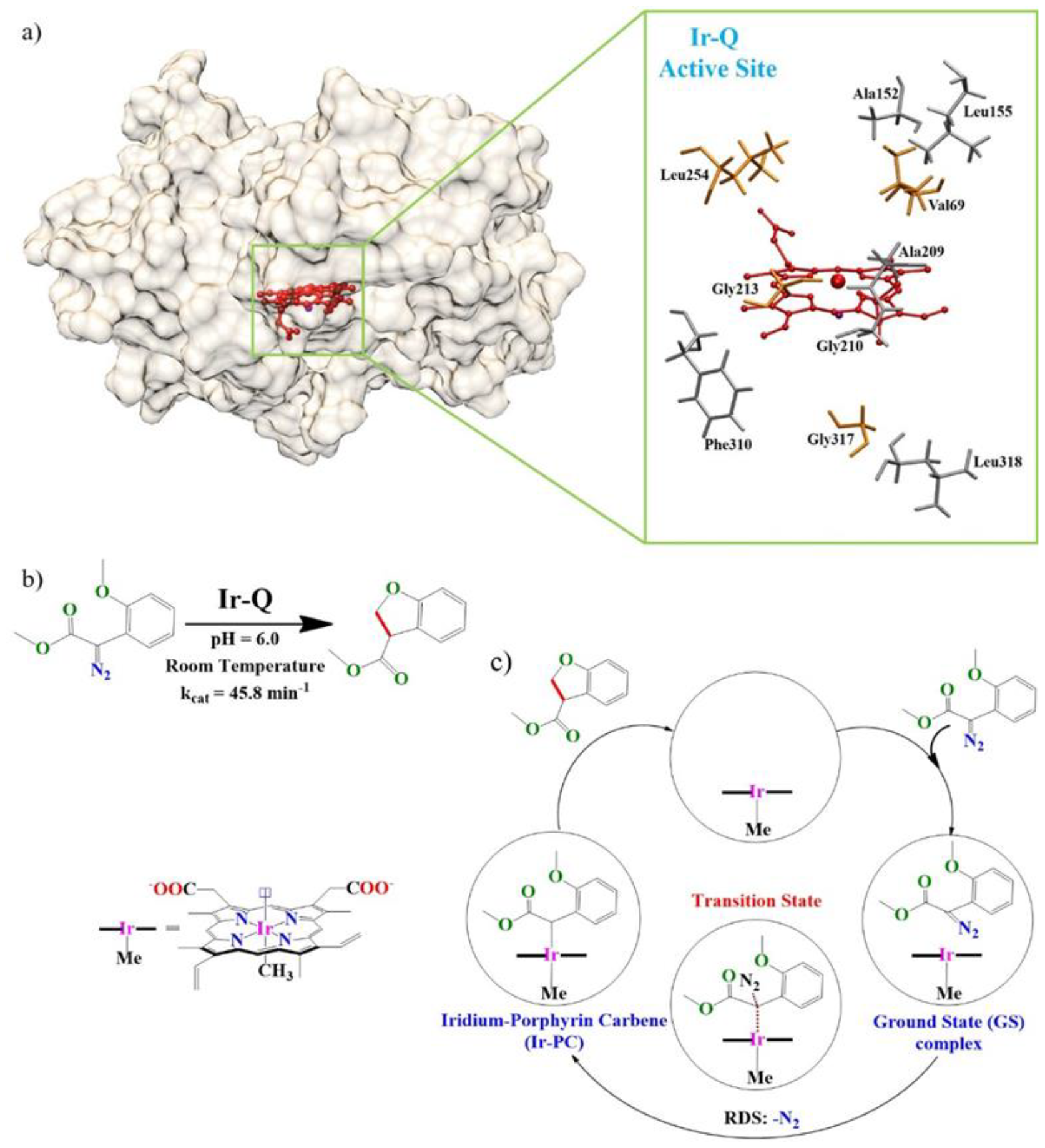
Schematic representations of (a) the most efficient quadruple mutant, **Ir-Q** artificial metalloenzyme with active site residues shown in the enlarged region. The red color denotes the Ir(Me)-porphyrin moiety with the orange color denotes the mutated active site residues and grey as other active side residues. (b) The chemical reaction catalyzed by **Ir-Q**.(7) (c) Proposed mechanism of the carbene insertion reaction catalyzed by **Ir-Q** based on reference (8). The black circle represents the protein scaffold. RDS, rate determining step.

There has been impressive progress in understanding the underlying rationale behind the directed evolution produced efficient artificial enzymes by using both computational(9) and experimental methods.(10)Arnold and coworkers have isolated and characterized Iron Porphyrin Carbene (**IPC**) intermediate inside a cytochrome c transferase and provide invaluable mechanistic insights related to directed evolution mutations.(11)They also reported isolation of a metallo-aryl-carbene intermediate in a Protoglobin, however no rationale for transition state stabilization is given.(12)In another recent study, Hilvert, Kern and coworkers has shown how directed evolution gradually altered the conformational ensemble of the protein scaffold to populate a narrow, highly active conformational ensemble and accelerate an elementary proton transfer reaction enormously.(13)

The directed evolution method, a top-down approach, has made so far the most successful advancement in the field of protein catalysis/enzyme design.(4, 14, 15) Although powerful, directed evolution is costly and time consuming which restricts the potential impact of biocatalysis on many industrial processes.(3, 16) Improving enzymes design protocols requires thorough understanding of the activity enhancements brought about by directed evolution.(3)It has been prescribed that information about the reaction energetics (stability of the transition state) is in principle a more reasonable direction to guide directed evolution.(17, 18) Motivated by the above exciting progresses, we embark on a journey to decipher the origin of natural-like high catalytic rate enhancement of the artificial enzyme **Ir-Q** in comparison to the single (**Ir-S**, C317G) and double (**Ir-D**, C317G, T213G) mutants using large-scale molecular dynamics simulations and rigorous quantum chemical calculations. Our results reveal in tandem with the previous works(13, 19)about how the directed evolution precisely position the cofactor-substrate in an unusual orientation within a reshaped active site that emerged during evolution, to attain maximum transition state stabilization to achieve enormous rate acceleration. While the electrostatic model is widely established, this work set a new design paradigm for improving enzyme design protocols, as laid down in recent reviews. (1, 3, 20)

## Results and Discussion

### Decoding the catalytically active conformation in the most efficient artificial metalloenzyme, Ir-Q

There were no crystal structures available for the most efficient quadruple mutant, **Ir-Q** as well as for the single (**Ir-S**) and double (**Ir-D**) mutants, which has precluded a mechanistic understanding of this highly efficient artificial enzyme. However, Hartwig’s recent report on a closely related **Ir-ArM** has provided us with crucial mechanistic details.(8) Based on these evidences, the catalytic mechanism of carbene insertion in **Ir-Q** is proposed as follows (Fig. 1c): (i) formation of a non-covalent GS (ground state) complex between the holoenzyme and the substrate (diazoester); (ii) a rate determining formation of **Ir-PC** intermediate due to its close resemblance to the **Ir-ArM** enzyme followed by asynchronous carbene insertion into the C−H bond. (21) Herein understanding of the RDS formation of **Ir-PC** inside the protein scaffold is pivotal to explain its remarkable catalytic efficiency. In order to map the conformational space, we subjected the **Ir-Q** mutant bound with diazoester substrate to extensive molecular dynamics (MD) simulations with three replica starting structures as follows. Two replicas of **Ir-Q** are generated with one (A) from the **Ir-ArM** crystal structure (PDB: 7UOR) and the other (B) from the wild type (WT) P450 enzyme CYP119 crystal structure (PDB: 1IO7) by appropriate mutations and diazoester docking in order to ascertain its actual conformation. Replica C is further generated from equilibrated trajectory of **Ir-ArM** crystal structure. Since the three replicas do not show any significant deviations, the results are discussed with respect to replica A and B only. We also generated the single (**Ir-S**, C317G) and double (**Ir-D**, C317G, T213G) mutants from the WT CYP119 crystal structure to capture the prevalent conformation ensemble in it.

The simulation results indicate minor changes in the global conformational dynamics of **Ir-Q** (Fig. 2), as was also shown in the Hartwig and coworker’s recent paper.(8)However, residue-based Root Mean Square Fluctuations (RMSF) values suggests that there is indeed a reorganization in the fluctuations of the active site residues in **Ir-Q** (SI Appendix, Fig. S2). Especially noteworthy is the reduced fluctuations of the Val69 and Leu254 residues in **Ir-Q** compared to that in **Ir-S** and **Ir-D** mutants. Indeed, Hartwig and coworkers indicated about the change in the shape of the active site in the closely related **Ir-ArM** as well as in the isolated aryl-IPC intermediate in Protglobin by Arnold and coworkers.(12)Hence, the vicinity of the active site in **Ir-Q** is probed in more details to decipher the origin of its extraordinary catalytic efficiency.

**Fig. 2.**
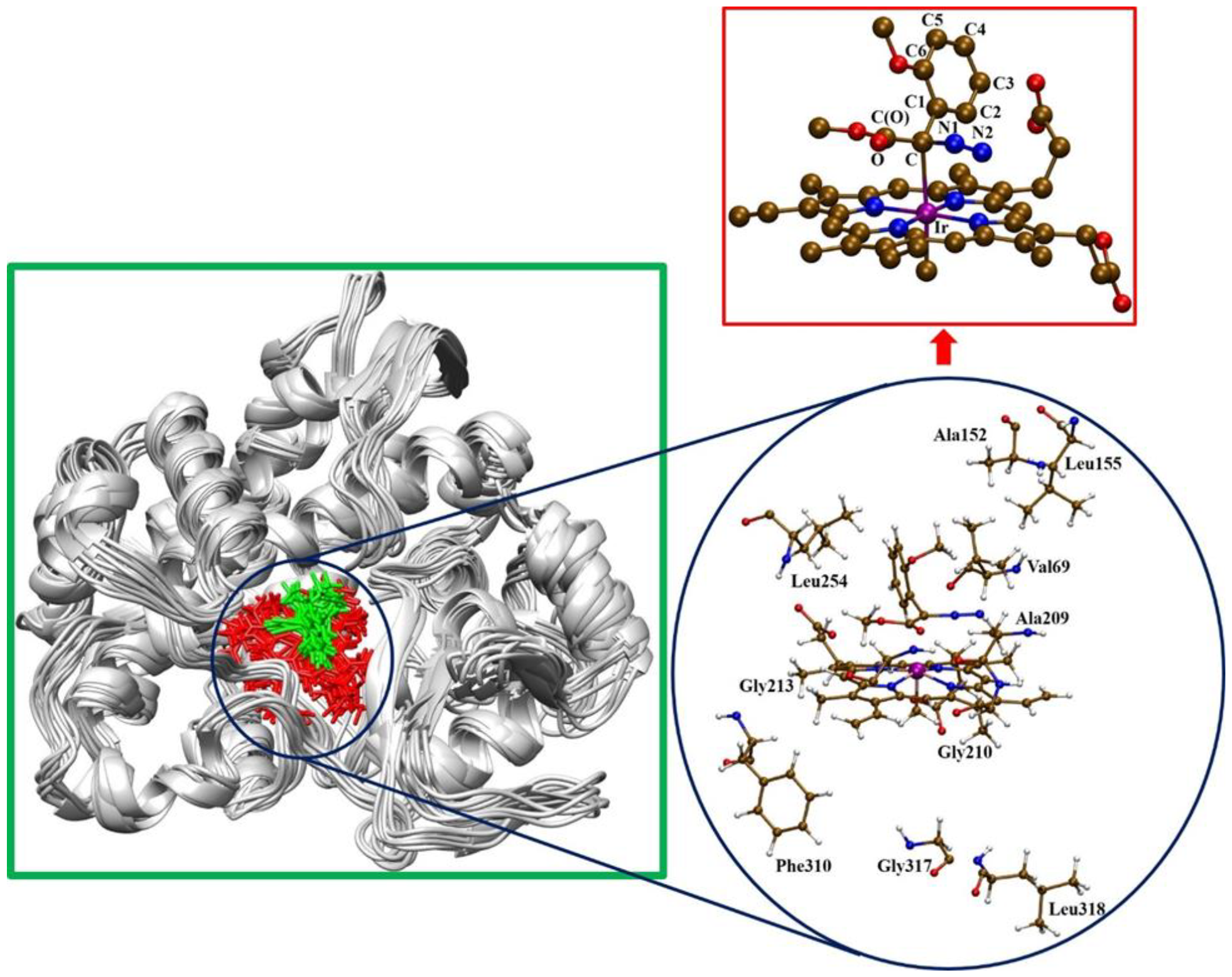
Overlay of 10 representative snapshots obtained from 500ns of classical molecular dynamics simulations of diazoester bound **Ir-Q** (A). The cofactor is represented in red and the substrate in green. (Inset) Illustration of the diazoester bound Ir-porphyrin cofactor with labeled active site residues and key atoms (top inset).

It is worthwhile to analyze the effect of these fluctuations in the active site residue dynamics on the Ir-porphyrin substrate (Ground State, GS) complex from simulation trajectory of A, B and C replicas of **Ir-Q**. The relative orientation of the substrate bound to the Ir-porphyrin cofactor is described along the simulation trajectory by two different dihedral angles of the **GS** complex; dihedral-1, ∠C2-C1-C-N1 and dihedral-2, ∠C2-C1-C-Ir (Fig. 2) in order to monitor the conformational variations. Notably the two dihedral angles are restrained in the A, B and C replicas of **Ir-Q** (SI Appendix, Fig. S3 and S4), in sharp contrast to the fluctuations observed in the analogous dihedrals during a similar simulation trajectory of the free Ir-cofactor complex in water (SI Appendix, Fig. S5). Most interestingly, two well-defined conformations have been characterized based on the orientation of the aryl (Ar) substituent of the substrate with respect to plane of the Ir-porphyrin ring in **Ir-Q** and other mutants as follows. One is the **Ir-Q**_planar_ conformation where the Ar group and the Ir-porphyrin ring are more or less coplanar (dihedral-1 ∼ 0° and dihedral-2 ∼ 90°) and the other one is **Ir-Q**_perpendicular_ where the two moieties are in a near or fully perpendicular orientation (dihedral 1 ∼ 90° and dihedral 2 ∼ 0°) (Fig. 3). The simulation data reveals a strong preference for the **Ir-Q**_perpendicular_ conformation ensemble (Fig. 3b) in all the three replicas of **Ir-Q**. On the contrary, the **Ir-Q**_planar_ conformation ensemble is found to be significantly populated in the **Ir-S** and **Ir-D** mutants.

**Fig. 3.**
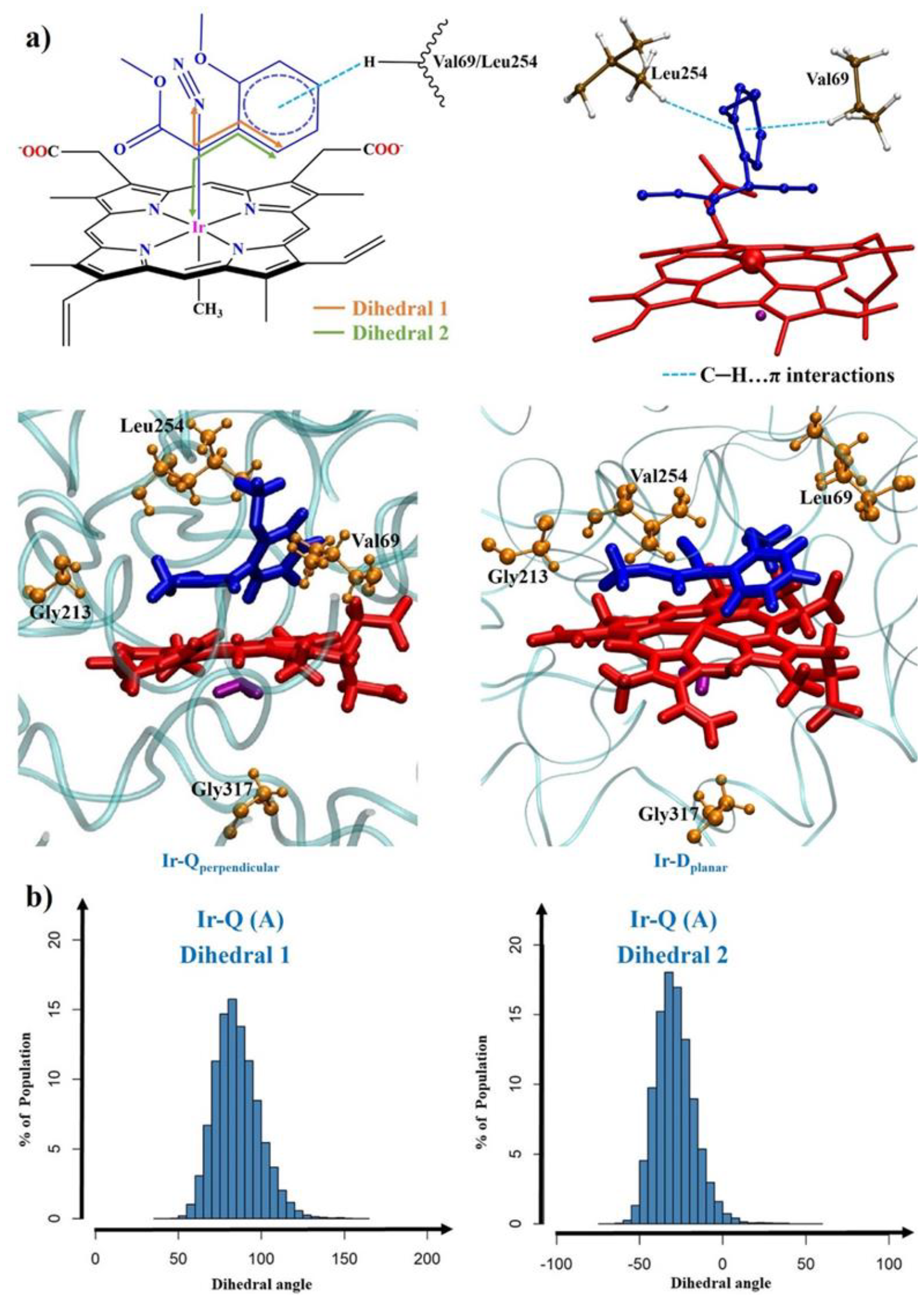
a) Top: Illustration of dihedral angles monitored during the MD simulation trajectory of the GS complex. Bottom: Representation of the two major types of conformations of the cofactor-substrate complex illustrating relative orientation of Ar group of the substrate (blue color) and the Ir-porphyrin ring (red color) [left side = **Ir-Q**_perpendicular_ conformation and right side = **Ir-D**_planar_ conformation]. Beside: More ordered Val69 and Leu254 residues emerged during evolution helps to populate the **Ir-Q**_perpendicular_ conformation. H atoms are omitted for clarity in the cofactor-substrate. b) Histogram population analysis of conformational states in the simulations trajectory of **Ir-Q** (A) based on the values of dihedral angles 1 and 2.

The distance between a π electron rich aryl C atom and a H atom of the Val69 sidechain in **Ir-Q**_perpendicular_ falls majorly between 3.0 to 4.0 Å (SI Appendix, Fig. S6a), which is conducive for a C−H…π interaction as is also proven by NCI (Non-Covalent interaction) analysis plot (SI Appendix, Fig. S7).(22)(23) Similar kind of NCI interaction is also possible with the V254L residue (SI Appendix, Fig. S6b). Interestingly, this restrained conformation is not observed in the trajectory of the **Ir-S** and **Ir-D** mutants, which may be ascribed to the more flexible Leu69 and Val254 residues. The reduced fluctuations of the Val69V and Leu254 residues along with the C− H…π interactions engender the Ir-porphyrin-substrate (cofactor-substrate) complex in a particular **Ir-Q**_**perpendicular**_ conformation. There are evidences of non-covalent interaction (e.g., π-π) playing a decisive role in the active configuration and reactivity of a Ir-carbene species in the free state as well as in the IPC intermediate inside an enzyme.(12, 24)While the C317G mutation accounts for accommodating the Ir(Me) moiety, T213G mutation envelope and help to orient the ester part of diazoester by hydrophobic interactions. These results are in excellent agreement with the reported **Ir-ArM** crystal structure.(8) However, our study clearly shows the role of the L69V and L254V mutations in attaining a well-defined **Ir-Q**_perpendicular_ conformation, which is different from that in **Ir-ArM**.

### Reproducing the experimental activation barrier height from ‘catalytically active conformation’, Ir-Q_perpendicular_

To probe the effect of this restrained conformation on the observed reactivity of the **Ir-Q** mutant, we have employed a quantum mechanical (QM) cluster model approach. In the cluster approach, a relatively small part of the enzyme around the active site are truncated at their α-carbons, that are fixed during optimization to preserve the constraints imposed by the protein’s secondary and tertiary structure, and is selected as a model for full QM calculations.(25)Siegbahn and coworkers along with others have pioneered the use of cluster model approach in investigating metalloenzymes, which provide accurate results as per any other computational methods.(26–28) Warshel has also approved this approach for metalloenzymes, especially in the context of less electrostatic influence on the active site from the global protein structure.(18)Herein, we have created two separate cluster models (model 1 and 2) which includes the Ir-porphyrin, substrate (diazoester) and the first and second sphere residues (69, 152, 155, 205, 209, 213, 254, 310, 317 and 318, see blue circled in Fig. 2) by considering the **Ir-Q**_perpendicular_ orientation from the snapshots of the simulation trajectories of **Ir-Q** (**A**) and **Ir-Q** (**B**) respectively to calculate the activation barrier of the rate determining step of **Ir-PC** intermediate formation. The reaction energy profile (Fig. 4) is shown in the singlet state as GS complex **1** is significantly lower (41.4 kcal/mol) than its triplet state. Since the **Ir-PC** formation step is calculated to be higher (14.5 kcal/mol) in energy than the carbene insertion step (1.9 kcal/mol), we focused on the RDS to explain the observed catalytic rate enhancement.

**Fig. 4.**
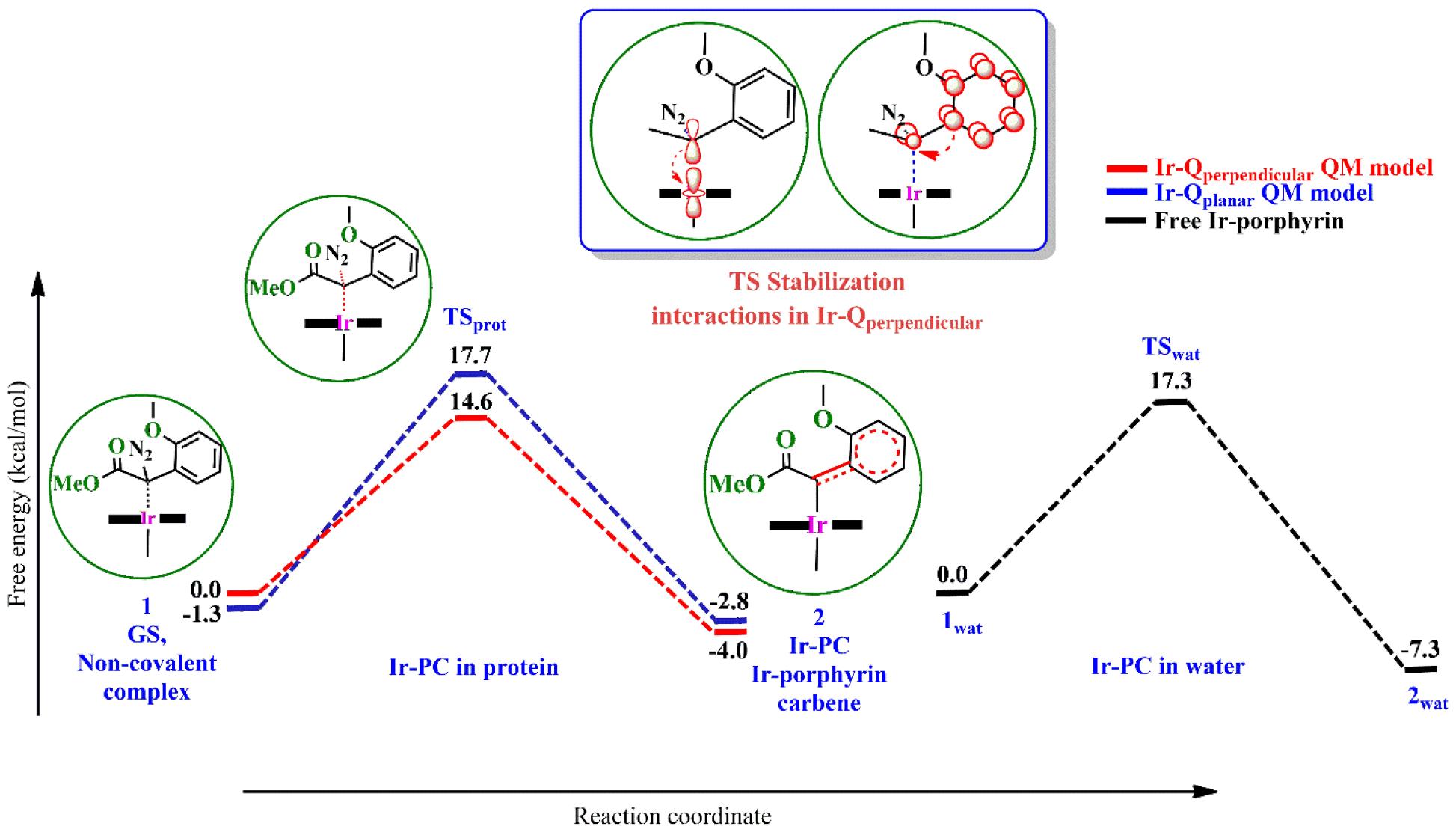
Reaction energy profiles of the RDS step for **Ir-PC** intermediate formation obtained using quantum chemical cluster models of **Ir-Q**_**perpendicular**_ and **Ir-Q**_**planar**_ conformations and the free Ir-cofactor complex in water at the PCM (ε=4.0/80.1)/ B3LYP-D3/6-311G**, LANL2DZ (Ir) // B3LYP/6-31G*, LANL2DZ (Ir) level of theory. (Inset) The preferential stabilization of the transition state in **Ir-Q**_**perpendicular**_ by backdonation of Ar π-electrons into the carbene p orbital. The green circle around the Ir-cofactor-complex represents the first and second sphere amino acid residues of the protein considered in the quantum chemical cluster model.

Our QM study shows that the calculated activation barrier (**ΔG**^**#**^) for the RDS, i.e. the **Ir-PC** formation/nitrogen loss step in **Ir-Q** is 14.6 kcal/mol. Similar activation barrier, ΔG^#^ **=**14.5 kcal/mol is also obtained from model 2. Most importantly, it agrees well with the experimentally observed barrier of 12.8 kcal/mol,(7)obtained from the k_cat_ value (Table 1).(7)Indeed, there is a lowering of the activation barrier for the formation of the **Ir-PC** intermediate inside the protein scaffold of **Ir-Q** compared to that of free Ir-cofactor complex in water (**ΔG**^**#**^ **=** 17.3 kcal/mol, Fig. 4 and Table 1). The difference in the activation energy for formation of the **Ir-PC** between the water and protein scaffold is 2.7 kcal/mol, which further explains the observed 20 times faster catalysis in **Ir-Q** than by the free Ir-cofactor.(7) The fact that the two quantum chemical cluster models generated from two distinct replicas result in a similar activation barrier for the **Ir-PC** intermediate formation reinforces the accuracy of our large-scale simulation for identifying the catalytically active conformation in directed evolution produced enzymes.

**Table 1.** Calculated and experimental activation energy barrier of **Ir-PC** formation step for two conformations of **Ir-Q** using quantum chemical cluster models and of the free Ir-porphyrin complex in water. Computed values are reported at the PCM (ε=4.0/80.1)/ B3LYP-D3/6-311G**, LANL2DZ (Ir) // B3LYP/6-31G*, LANL2DZ (Ir) level of theory. Energy values are in kcal/mol.

| Quantum chemical cluster models | Activation energy barrier ($\Delta E^\ddagger_{\text{calc.}}$) | Free energy of activation, ($\Delta G^\ddagger_{\text{calc.}}$) | $\Delta G^\ddagger_{\text{expt}}$ (from observed $k_{\text{cat}}$) |
| --- | --- | --- | --- |
| **Ir-Q<sub>perpendicular</sub>** | 14.6 | 14.6 | 12.8 |
| **Ir-Q<sub>planar</sub>** | 23.7 | 19.0 |  |
| **Free Ir-porphyrin complex** | 20.1 | 17.3 |  |

Considering the existence of **Ir-Q**_planar_ conformation in **Ir-S** and **Ir-D** mutants, we also explore the effect of this specific Ir-porphyrin-substrate orientation inside the protein scaffold by adjusting the dihedrals to generate a cluster model resembling **Ir-D**_planar_ conformation (Fig. 3). The calculated activation barrier for **Ir-PC** formation in **Ir-Q**_planar_ is found to be 19.0 kcal/mol, which is significantly higher than the **Ir-Q**_perpendicular_ conformation (Table 1). Since **Ir-Q**_perpendicular_ and **Ir-Q**_planar_ conformations has a similar protein environment, it can be conservatively said that the electrostatic effect of the protein may not have any significant effect in the reduction of the free energy activation barriers. This, however, appears understandable considering the hydrophobic nature of the active site and the Ir-cofactor-substrate complex. The catalytically active **Ir-Q**_perpendicular_ conformation, which only reproduces the experimental observed activation barrier, has been established by our rigorous large-scale simulations from three independent replicas derived from two different crystal structures. The productive **Ir-Q**_perpendicular_ conformation in the most efficient enzyme thus belongs to the repertoire of ‘catalytically active’ conformational states on the reactivity of enzymes.(29)

### Underlying factors for catalytic rate enhancement

To understand the underlying factors behind the relative energies of the transition states (TS), we employed Molecular Orbital (MO) and Natural Bond Orbital (NBO) analysis to decipher the nature of bonding in the Ir-porphyrin-substrate complex. All the TS involve a concerted cleavage of the C−N bond and development of an Ir−C bond along with a developing electrophilic carbene C. The calculated Ir−C bond lengths in **Ir-Q**_**perpendicular**_cluster model varies from 3.64Å to 2.399Å from GS to TS indicative of Ir−C bond formation (standard Ir−C = 2.055Å), as is also supported by MO bonding and NBO analysis (SI Appendix, Fig. S9).

Especially unusual is the stabilization of the developing vacant p orbital at the carbene C center due to the leaving N_2_ group in the TS by backdonation of π electrons from the Ar ring owing to its perpendicular orientation in **Ir-Q**_**perpendicular**_ fostered by C−H…π interactions of more ordered Val69 and Leu254 sidechains. This Ar… C_carbene_ conjugation cannot be effectively operational in the GS which provides the differential TS stabilization. It is further evident from the relatively shorter C−C1 bond lengths (1.480Å, Table 2) as well as MOs, second order NBO stabilization energy and NICS calculations. This synergistic C…Ir dative σ bonding and Ar…C π backdonation, which is not present significantly in the TSs of Ir-Q_**planar**_ and the free Ir-porphyrin complex, essentially provide preferential stabilization to the TS in **Ir-Q**_**perpendicular**_ to reduce the activation free energy barrier for the RDS step. Unlike the usual bonding description in metal-carbene complexes, we argue that the Iridium porphyrin carbene (**Ir-PC)** presents an unprecedented bonding situation in a metal-carbene intermediate with synergistic C…Ir dative σ bonding along with a π donor Ar substituent inside a protein scaffold.(30, 31)

**Table 2.**
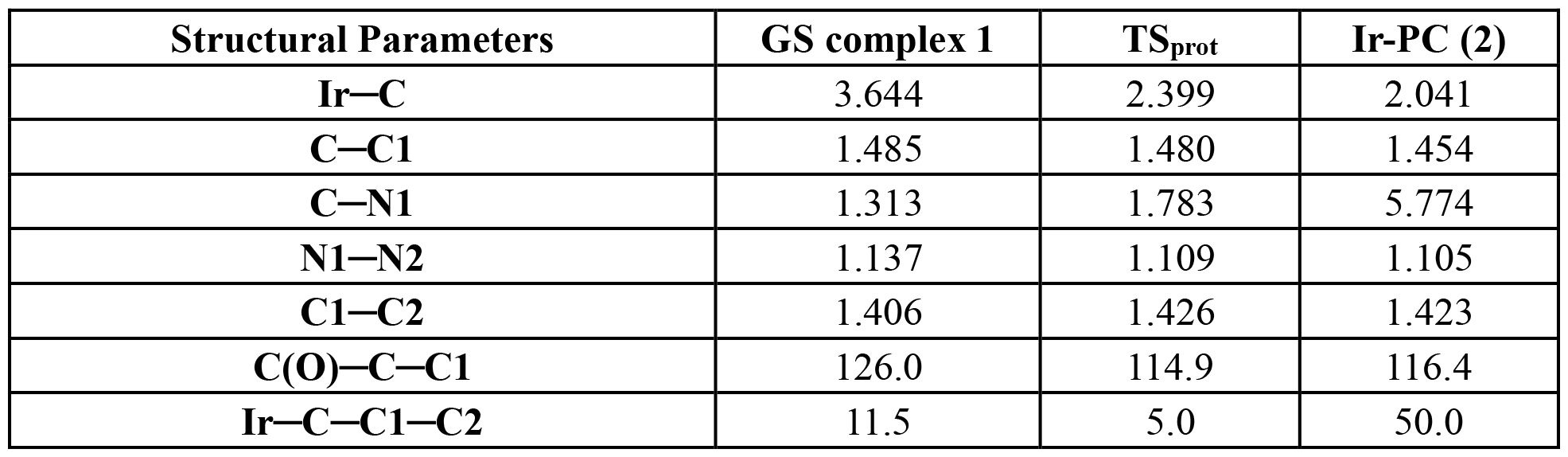
Important structural parameters of optimized structures of the enzyme-substrate, transition state and intermediate cluster models of **Ir-Q**_perpendicular_ at the B3LYP/6-31G*, LANL2DZ (Ir) level of theory (see Fig. 4). All bond lengths are in Å and angles are in degrees.

| Structural Parameters | GS complex 1 | TS<sub>prot</sub> | Ir-PC (2) |
| --- | --- | --- | --- |
| Ir–C | 3.644 | 2.399 | 2.041 |
| C–C1 | 1.485 | 1.480 | 1.454 |
| C–N1 | 1.313 | 1.783 | 5.774 |
| N1–N2 | 1.137 | 1.109 | 1.105 |
| C1–C2 | 1.406 | 1.426 | 1.423 |
| C(O)–C–C1 | 126.0 | 114.9 | 116.4 |
| Ir–C–C1–C2 | 11.5 | 5.0 | 50.0 |

Our study thus shows how the protein scaffold preorganized the **GS** complex in **Ir-Q**_**perpendicular**_ to avail of the maximum TS stabilization to exhibit high catalytic rate acceleration. Mulholland and coworkers had previously reported on selective TS stabilization in Kemp eliminase to account for its high catalytic efficiency.(32)The end-on orientation of the IPC intermediates inside cytochrome c tranferase is held on by stabilizing hydrophobic interactions.(11) In contrast, the perpendicular orientation of the Ar ring and Ir-porphyrin ring in **Ir-Q**_**perpendicular**_ is held on by C−H…π interactions from more ordered mutated L69V and V254L residues. Encouragingly, this kind of restrained orientation of the Ar group in an **Ir-PC** intermediate by non-covalent interactions gets good support from recent isolation of an aryl Iron-carbene intermediate inside Protoglobin by the same group,(12) which, however, did not reveal the underlying TS stabilizing interactions like that in this study. Thus, this preferred conformation is the main reason to provide preferential TS stabilization in **Ir-Q** compared to the other conformations and the free Ir-porphyrin complex.

## Conclusions

Our computational study reveals how directed evolution mutations evolve the conformational states of **Ir-Q** to generate a catalytically productive active site in the **Ir-Q**_perpendicular_ conformation ensemble, which is primarily responsible for the natural-like rate enhancement of the artificial metalloenzyme. It reveals the effect of L69V and V254L directed evolution mutations in atomistic details in lowering the activation energy barrier for the RDS step of **Ir-PC** formation to render the remarkable catalytic efficiency. The catalytically active conformation features an unprecedented C…Ir bonding and Ar…C π backdonation that preferentially stabilizes TS to exhibit high catalytic rate enhancement. Moreover, it shows how mutations acquired during directed evolution precisely position the cofactor-substrate complex in a productive orientation inside the protein to achieve effective transition state stabilization compared to the ground state. We also conjecture that **Ir-Q** with a more electron-rich substrate Ar substituent should surpass even the natural enzymes in terms of catalytic rate enhancement. The catalytic factors revealed in this study would help to design a ‘complex theozymes’(3) to enable a more effective discrimination of the transition state to the ground state inside a protein scaffold. Moreover, this elegant strategy could be utilized to stabilize such unusual ‘complex theozymes’ to improve enzyme design methods in conjunction with data-driven approaches(33, 34) in view of increasing computational power. Our study thus reports a new design paradigm apart from the electrostatic model towards realization of fully programmable protein catalysis.

## Methods

Molecular Dynamics simulations were performed using the GPU code of the NAMD package.(35) For the classical MD simulations, the CHARMM force field was utilized. (36)We utilized the two reported crystal structures (PDB ID: 7UOR and 1IO7) for generating the initial coordinates for the simulations. The time step for the MD simulations was taken as 1 fs. The simulations were run for 500ns. All QM calculations were carried out with the DFT method implemented in the Gaussian 16 program.(37) The B3LYP functional along with the LANL2DZ basis set with an effective core potential (ECP) for Ir and standard split valence Pople’s basis set 6-31G* for all other atoms were used for the optimizations,(38–40)along with the D3 version of Grimme’s dispersion correction.(41, 42). We created the cluster models(43)from appropriate snapshots from the molecular dynamics simulations for quantum chemical (QM) calculations.

## Supporting information

Fig. S1. Protein backbone (Calpha) RMSD plots for three replicas of Ir-Q from the MD simulation trajectory.

Fig. S2. RMSF plots for the protein backbone (Calpha) atoms for 500ns of the simulation shown for the WT protein CYP119, Ir-S, Ir-D and Ir-Q

Fig. S3. Variation of dihedral 1 during the course of the simulation for different replicas of Ir-Q.

Fig. S4 Variation of dihedral 2 during the course of the simulation for different replicas of Ir-Q.

Fig. S5. Illustration of a) a snapshot of the free Ir-porphyrin-substrate complex in water from simulation trajectory and b) variations of dihedral 1

Fig. S6. C H pi interaction distance (top) in Ir-Q from a Aryl C atom of the substrate to H atoms of V69 & L254 sidechains

Fig. S7. Noncovalent interaction analysis plot illustrating the non-covalent interactions between a Aryl C atom and H atom of V69 sidechains

Fig. S8. Noncovalent interaction analysis plot illustrating the non-covalent interactions between an Aryl C atom and H atom of L254 sidechains

Fig. S9. MO Diagram of the critical Ir-Qperpendicular TS1 optimized structure demonstrating the stabilizing bonding interactions

## Data Availability

All data are available in the main text and SI.

## Acknowledgements

S. R. acknowledges support from Ramalingaswami Re-entry Fellowship (BT/RLF/Re-entry/39/2017), Department of Biotechnology, Govt. of India. The financial support from Saha Institute of Nuclear Physics, Kolkata is duly acknowledged. A.M. is thankful to SINP for providing a postdoctoral fellowship. We thank SINP computing facility for the computational resources.

## Classification

Biophysics and Computational Biology

