## Supplementary figures and images for "Non-Electrostatic Basis for an Artificial Metalloenzyme Catalysis"

### Fig. S1. Protein backbone (Calpha) RMSD plots for three replicas of Ir-Q from the MD simulation trajectory.

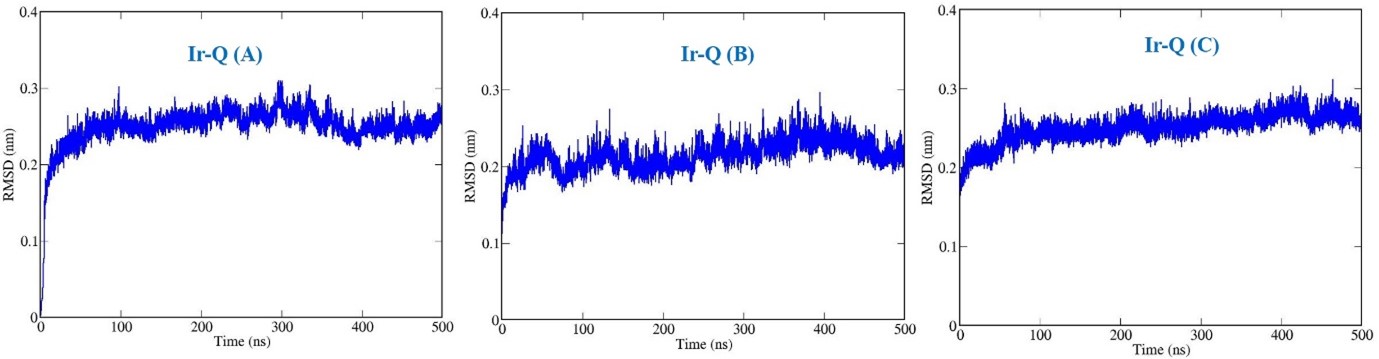

### Fig. S2. RMSF plots for the protein backbone (Calpha) atoms for 500ns of the simulation shown for the WT protein CYP119, Ir-S, Ir-D and Ir-Q

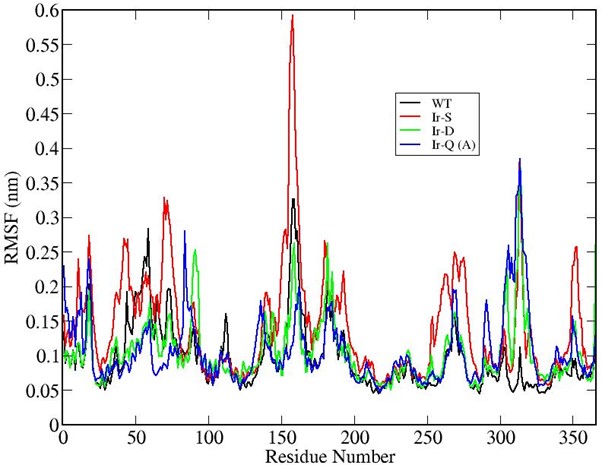

### Fig. S3. Variation of dihedral 1 during the course of the simulation for different replicas of Ir-Q.

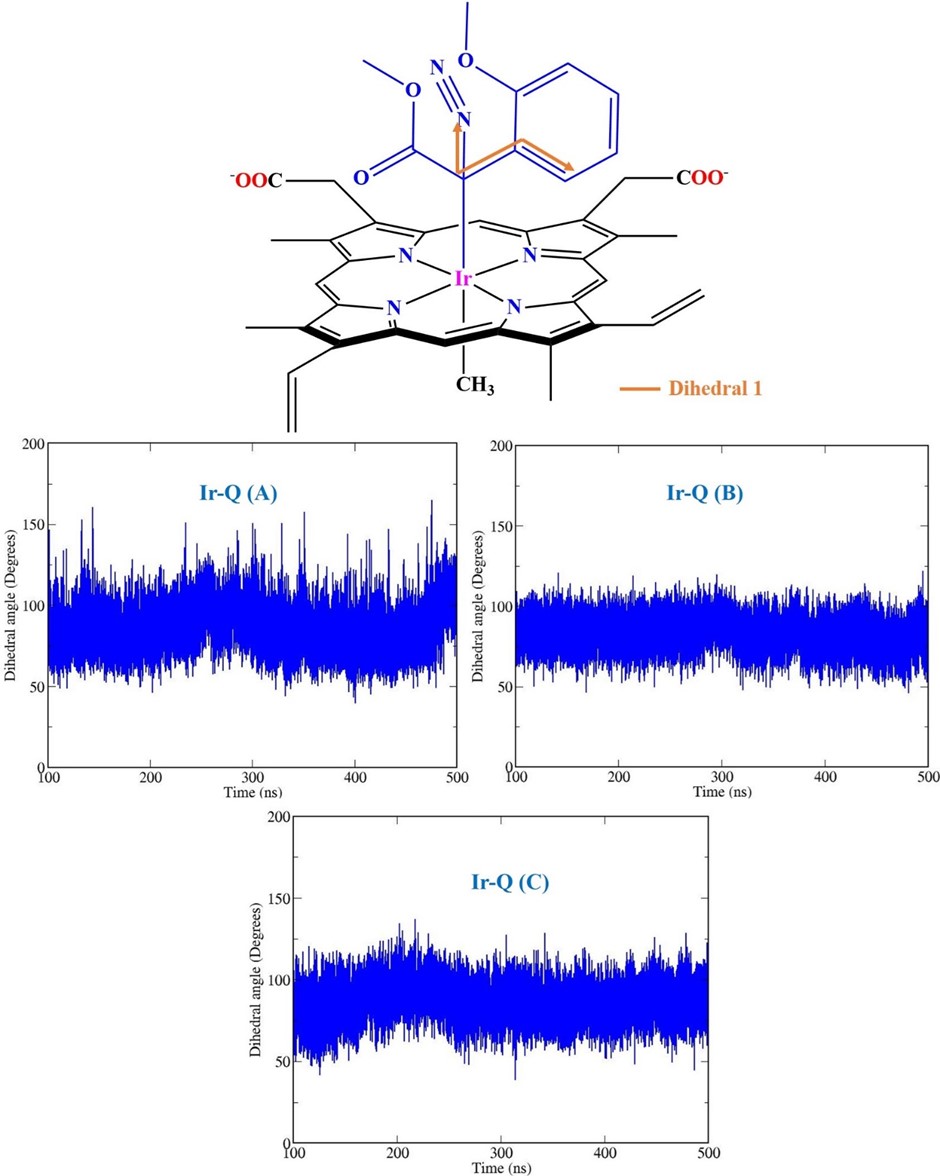

### Fig. S4 Variation of dihedral 2 during the course of the simulation for different replicas of Ir-Q.

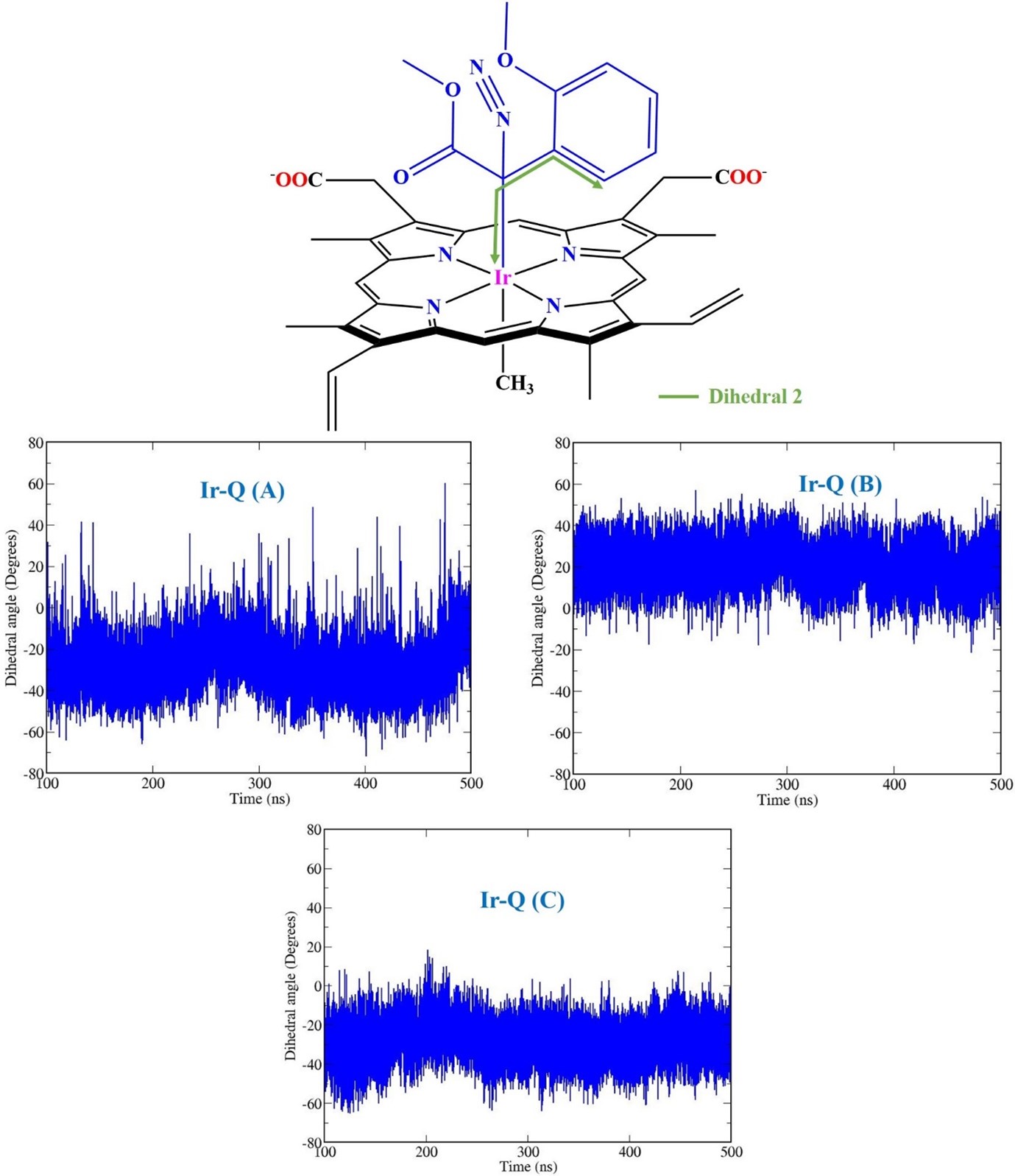

### Fig. S5. Illustration of a) a snapshot of the free Ir-porphyrin-substrate complex in water from simulation trajectory and b) variations of dihedral 1

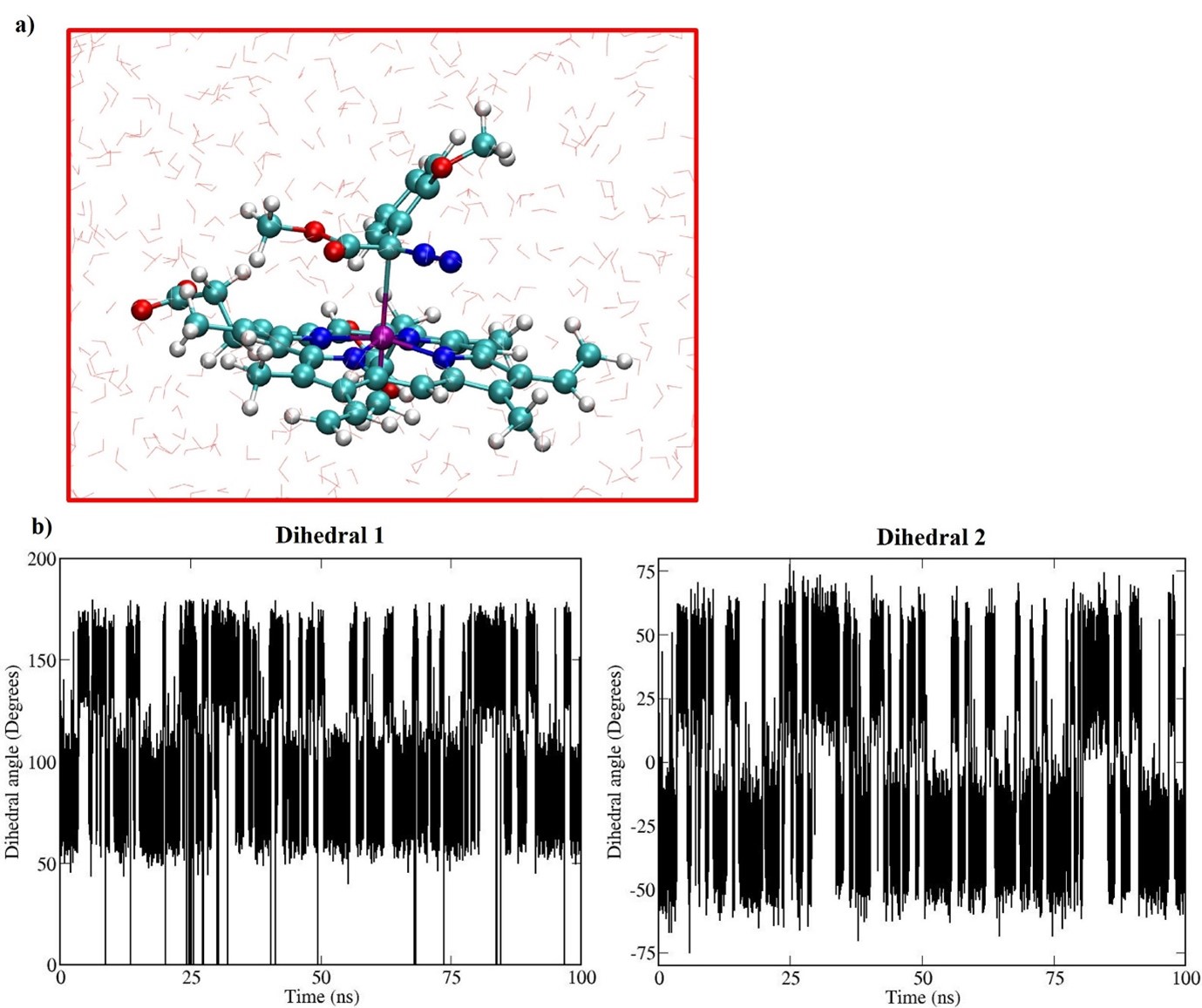

### Fig. S6. C H pi interaction distance (top) in Ir-Q from a Aryl C atom of the substrate to H atoms of V69 & L254 sidechains

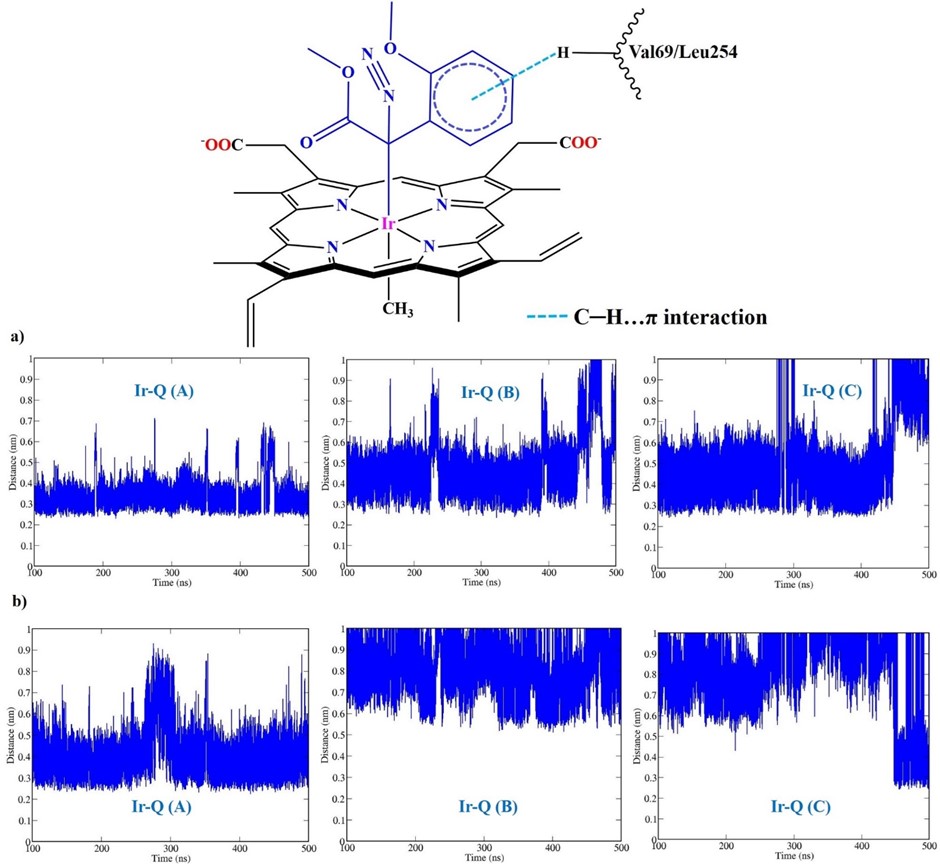

### Fig. S7. Noncovalent interaction analysis plot illustrating the non-covalent interactions between a Aryl C atom and H atom of V69 sidechains

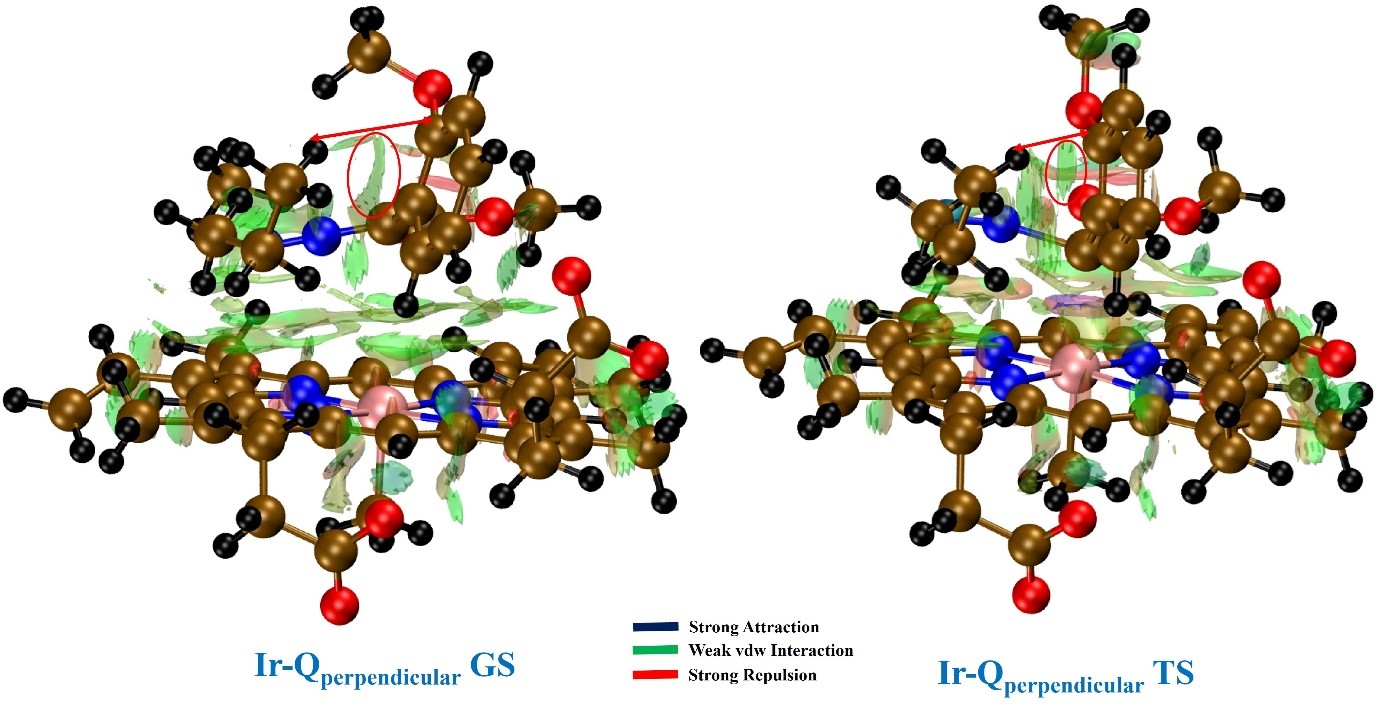

### Fig. S8. Noncovalent interaction analysis plot illustrating the non-covalent interactions between an Aryl C atom and H atom of L254 sidechains

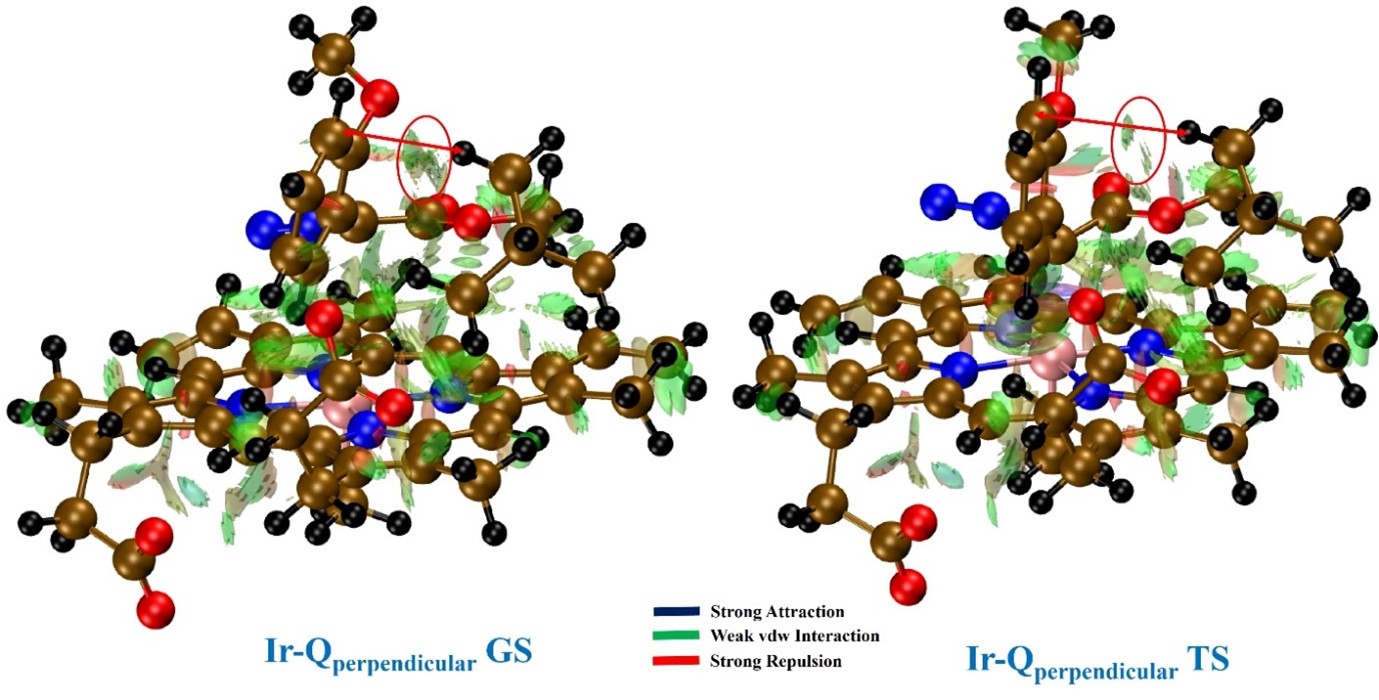

### Fig. S9. MO Diagram of the critical Ir-Qperpendicular TS1 optimized structure demonstrating the stabilizing bonding interactions

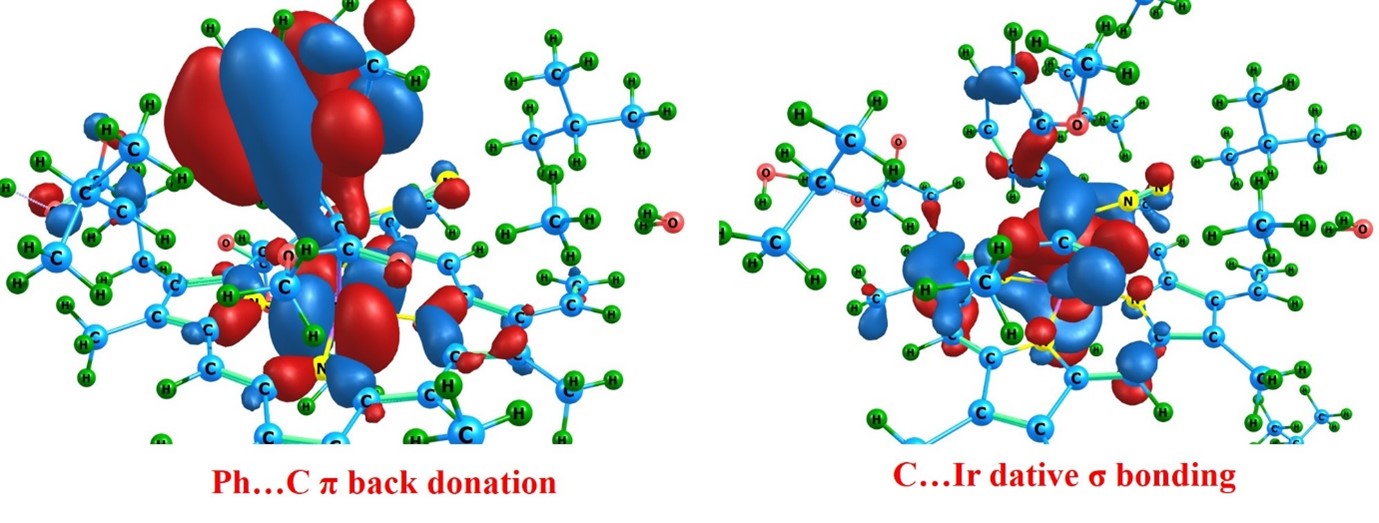
